# A Neural Microcircuit Model for a Scalable Scale-invariant Representation of Time

**DOI:** 10.1101/327387

**Authors:** Yue Liu, Zoran Tiganj, Michael E. Hasselmo, Marc W. Howard

## Abstract

Scale-invariant timing has been observed in a wide range of behavioral experiments. The firing properties of recently described *time cells* provide a possible neural substrate for scale-invariant behavior. Earlier neural circuit models do not produce scale-invariant neural sequences. In this paper we present a biologically detailed network model based on an earlier mathematical algorithm. The simulations incorporate exponentially decaying persistent firing maintained by the calcium-activated nonspecific (CAN) cationic current and a network structure given by the inverse Laplace transform to generate time cells with scale-invariant firing rates. This model provides the first biologically detailed neural circuit for generating scale-invariant time cells. The circuit that implements the inverse Laplace transform merely consists of off-center/on-surround receptive fields. Critically, rescaling temporal sequences can be accomplished simply via cortical gain control (changing the slope of the f-I curve).

## Introduction

### Behavioral evidence for a scale-invariant internal representation of time

Numerous behavioral experiments in humans and other animals suggest that time is represented in the brain in a scale-invariant fashion. For example, in interval timing experiments, the variability of the reproduced interval is proportional to the duration of the interval (Rakitin et al., 1998; Ivry & Hazeltine, 1995). The distributions of the response to different intervals are scale-invariant in that they overlap when rescaled by the duration of the interval, a phenomenon termed the scalar property (Gibbon, 1977).

Scale-invariance is also often observed in the associative learning rate in animal conditioning experiments. For instance, it has been shown that the number of trials needed for animals to develop a conditioned response increases when the reinforcement latency is increased and decreases when the intertrial interval is increased (Gallistel & Gibbon, 2000). Moreover, as long as the ratio between the intertrial interval and the reinforcement latency is fixed, the number of trials needed to develop a conditioned response is fixed, again indicating scale-invariance in the animal’s timing behavior.

Results from memory experiments also point to a scale-invariant representation of time. The classic power-law of forgetting (Wixted, 2004) indicates that a single mechanism may underlie both short and long term forgetting. In free recall, subjects are given a list of words and are asked to recall them in any order. The recency effect refers to the phenomenon that words from the end of a list are more easily recalled. This effect has been observed over a wide range of timescales, from fractions of seconds (Murdock & Okada, 1970) to several minutes (Glenberg et al., 1980; Howard, Youker, & Venkatadass, 2008), indicating that a single memory mechanism with a scale-invariant representation of time may serve under different timescales.

### Time cells in the brain

Behavioral scale-invariance requires that the neural system supporting behavior is also scale-invariant. Recent neurophysiological recordings in behaving animals show spiking activity at specific temporal intervals by individual neurons, referred to as *time cells*. These experimental data provide a possible neural substrate for timing behavior and various forms of memory (Howard, Shankar, Aue, & Criss, 2015).

Sequentially-activated time cells have been observed in a wide range of behavioral tasks and in many brain regions. Time cells were observed when an animal is performing delayed match to sample (MacDonald, Lepage, Eden, & Eichenbaum, 2011), delayed match to category (Tiganj, Cromer, Roy, Miller, & Howard, 2018), spatial alternation (Salz et al., 2016), or temporal discrimination tasks (Tiganj, Kim, Jung, & Howard, 2017). Time cells have been found in various parts of the brain including the hippocampus (MacDonald et al., 2011; Salz et al., 2016), prefrontal cortex (PFC) (Jin, Fujii, & Graybiel, 2009; Tiganj et al., 2017; Bolkan et al., 2017) and striatum (Adler et al., 2012; Mello, Soares, & Paton, 2015; Akhlaghpour et al., 2016). A recent study suggests that neurons in the amygdala are sequentially activated during the intertrial interval of a conditioning task (Taub, Stolero, Livneh, Shohat, & Paz, 2018).

Time cells exhibit phenomena that are suggestive of time-scale-invariance. The firing fields of time cells that fire later in the delay period are wider than the firing fields of time cells that fire earlier in the delay period (Figure 1). Moreover, the number density of time cells goes down with delay. Although there is not yet quantitative evidence that time cells are scale-invariant, these findings imply that the representation of the past is compressed (Howard, 2018) and are at least qualitatively consistent with a scale-invariant representation. If it turns out that sequentially-activated time cells support timing behavior, and if time cells are scale-invariant, then the neurophysiological mechanisms that endow time cells with scale-invariance are of critical importance in behavior. Scale-invariance of time cells is important because it provides the temporal basis for an animal to use the same set of mechanisms to integrate information and make decisions over different time scales. Because the natural world’s choice of scale is not known *a priori*, treating all scales equally is adaptive. It can be shown that a logarithmically spaced one-dimensional receptor optimally represent a function when the statistics of the stimulus function is unknown (Howard & Shankar, 2018). Just as in visual system the acuity decreases further away from the fovea and facilitates saccades, a scale-invariant representation of time where the temporal acuity decreases as we recede into the past would potentially facilitate retrieval of episodic memory (Howard, 2018).

### Chaining models are ill-suited to support scale-invariant sequential activation

A natural proposal for a model that generates sequentially activated neurons is to connect neurons sequentially in a one-dimensional chain. Although such chains can readily model sequentially-activated neurons, it is very difficult to make such chains scale-invariant.

For instance, Goldman (2009) proposed a feedforward network model for sustained persistent activity in a network in which a series of neurons modeled by leaky integrators are sequentially connected. In particular, the neurons all have the same decay time constant τ. By solving the dynamical equation it can be shown that their activations are given by the “time basis functions” 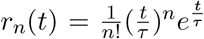. However this set of activations is not scale-invariant, as they are not of the same functional form. Figure 2 shows the actual and scaled neuronal activity in the chain. The rescaled neuronal activity becomes more concentrated for the neurons thatare activated later.

**Figure 1.**
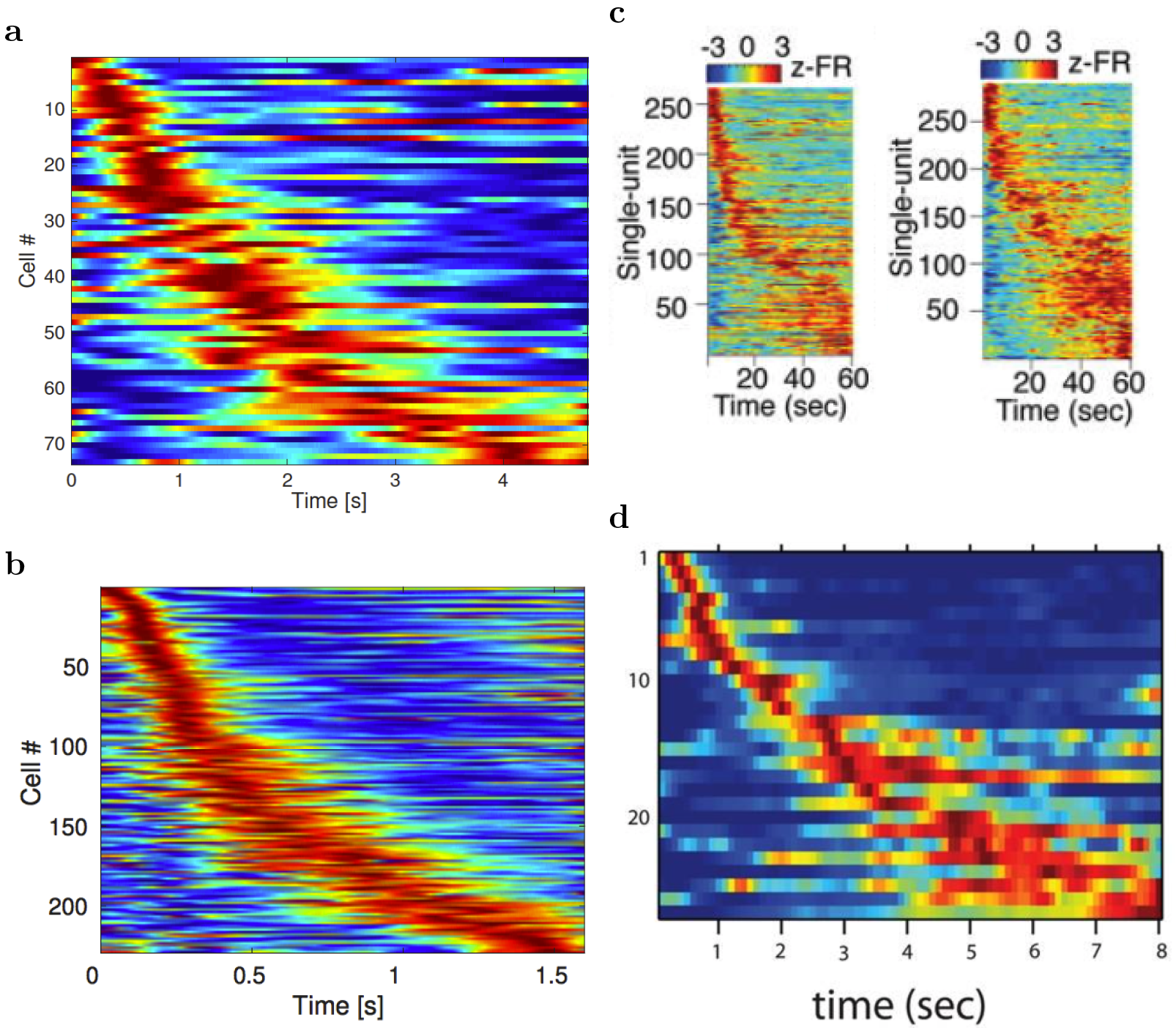
Sequentially activated neurons in the brain. Each row on each heatplot displays the normalized firing rate for one time cell. Red corresponds to high firing rate, while blue corresponds to low firing rate. The cells are sorted with respect to the median of the spike time in the delay interval. Two features related to temporal accuracy can be seen from examination of the heatmaps. First, time fields later in the delay are more broad than time fields earlier in the delay. This can be seen as the widening of the central ridge as the peak moves to the right. In addition the peak times of the time cells were not evenly distributed across the delay, with later time periods represented by fewer cells than early time periods. This can be seen in the curvature of the central ridge; a uniform distribution of time fields would manifest as a straight line. **a.**After Tiganj, et al., 2018. **b.** After Tiganj, et al., 2017. **c.** After Bolkan et al., 2017. **d.** After MacDonald et al., 2011.

This property ultimately arises from the Central Limit Theorem. Consider a chain of *N* neurons where every neuron is modeled by a same synaptic kernel *K*(*t*). That is, the activity of every neuron is the convolution of its synaptic kernel with the activity of the previous neuron.

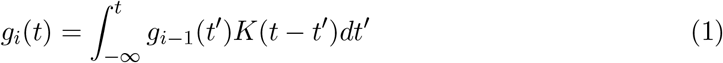

Since the functions *K(t)* and *g*_*i*_(*t*) are bounded, we can treat them as probability distributions up to a scale factor. Then the activity of the *i*th neuron *g*_*i*_ (*t*) is proportional to the probability distribution of the sum of the random variables described by *K(t)* and *g*_*i*–1_(*t*). Assuming the kernel *K* has mean *μ* and standard deviation σ, then by the Central Limit Theorem, for large *i*, *g*_*i*_(*t*) would have a Gaussian shape with mean *iμ* and standard deviation 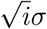. The coefficient of variation (CV) would scale as 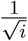. When rescaled, the neuronal activity for the neurons that get activated later will be more concentrated. This is indeed what is observed in Figure 2.

This logic is quite general. Systems that develop slow behavior from interactions among elements with a single characteristic time scale will show Central Limit Theorem scaling and thus not exhibit scale-invariance. This logic applies whether the kernel with a single characteristic time scale takes the form of a single time constant for the leaky integrators, a single time scale of synaptic transmission or a single time constant of a recurrent network. In order to construct a scale-invariant neural system, it is essential that it be endowed with a range of characteristic time scales.

Previous work (which we describe in detail below) has shown that one can build a scale-invariant memory using leaky integrators taking input in parallel, rather than in series as in the chaining model described above, if the integrators decay with a spectrum of time constants. This set of leaky integrators represents the Laplace transform of the past; approximately inverting the Laplace transform generates a set of units that activate sequentially and are scale-invariant (Shankar & Howard, 2013), much like neurophysiologically observed time cells (Howard et al., 2014). In this paper we develop a biologically-realistic minimal neural circuit to implement these equations. The set of leaky integrators with a spectrum of time constants is implemented using a previous computational model (Tiganj, Hasselmo, & Howard, 2015) that uses known single-unit properties of neurons in a variety of brain regions measured from slice physiology experiments (Egorov, Hamam, Fransén, Hasselmo, & Alonso, 2002; Fransén, Tahvildari, Egorov, Hasselmo, & Alonso, 2006; Navaroli, Zhao, Boguszewski, & Brown, 2011). We will find that the minimal circuit model to implement the inverse Laplace transform merely implements off-center/on-surround receptive fields. Moreover temporal rescaling of sequences can be accomplished by any mechanism that changes the slope of the f-I curve of the leaky integrators.

**Figure 2.**
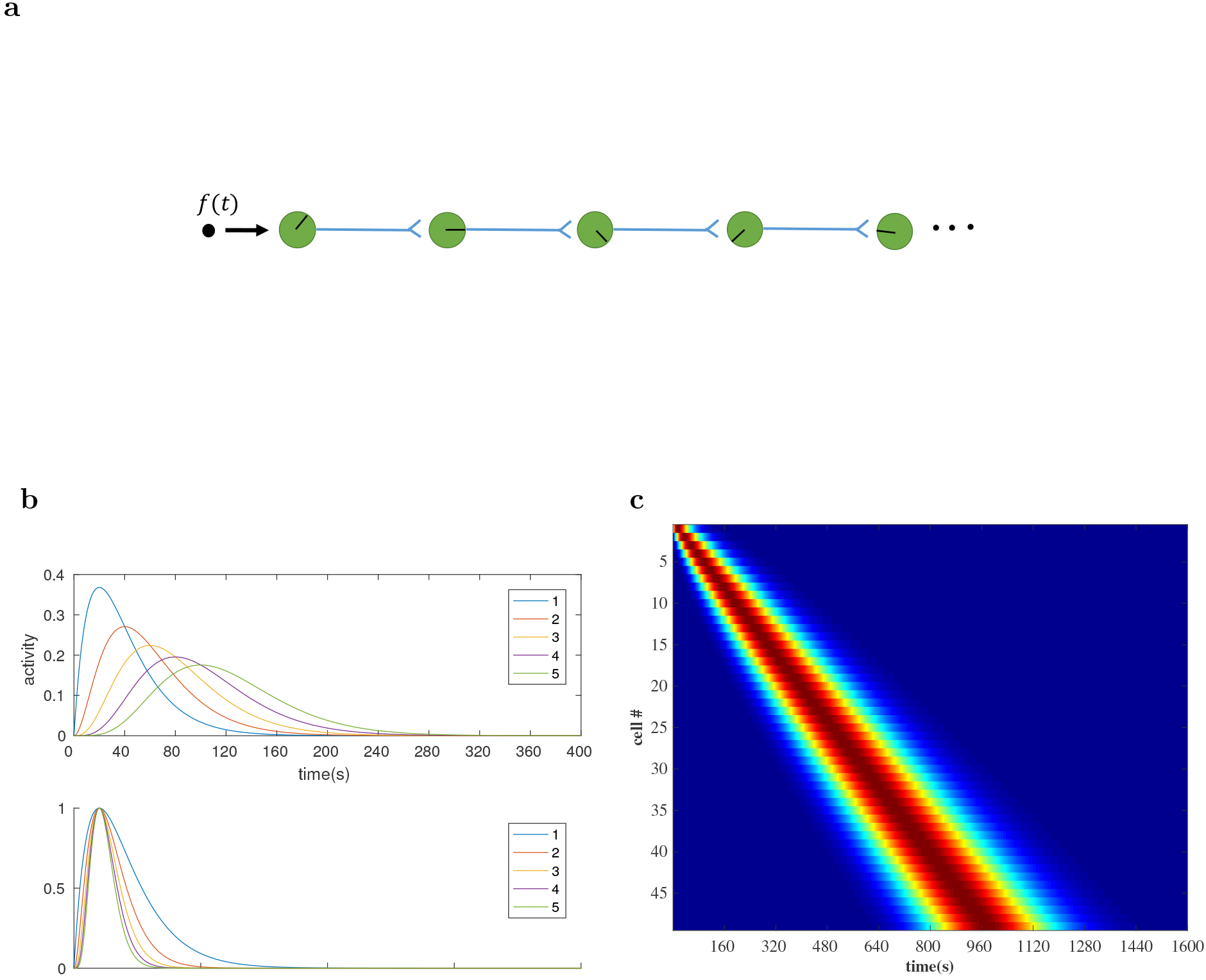
Simple chaining models produce time cells, but these time cells are not scale-invariant and have properties that differ from experimentally-observed time cells. **a:** A simple chain of units can give rise to sequentially-activated cells. The direction of the ‘clock hands’ within the neurons indicate the peak firing time of that neuron, **b.** Simulated “time basis functions” constructed from the chaining model described in Goldman (2009). **Top:** 5 successive time basis functions in the chaining model. They represent the neuronal activity for 5 successive nodes along the chain. We set τ = 20*s*. **Bottom:** The same time basis functions rescaled by the peak time along the x axis and by the maximum activity along the y axis. They deviate from each other systematically. It is clear that the neurons at different points along the chain do not obey scale-invariance. In particular, the activity of neurons that are activated later is more concentrated when rescaled. This can be shown in an asymptotic analysis using the Central Limit Theorem (see text), **c.** The heatmap for the activity of 50 neurons in the chaining model. The number of neurons coding later time is the same as the number coding for earlier times. This indicates that these neurons do not represent time in a scale-invariant way.

## Methods

Here we describe the mathematical framework for building a set of scale-invariant time cells. Following that, we describe a biologically-plausible instantiation of these equations.

### A mathematical approach for constructing a scale-invariant history

The neural circuit presented in this paper is built upon a mathematical framework that has been proposed to construct a representation of the recent past in a distributed, scale-invariant way (Shankar & Howard, 2012, 2013). This mathematical model has two layers of nodes. Nodes in the first layer integrate the input stimuli with an exponential kernel, equivalent to performing a Laplace transform on the stimuli. The activity of the nodes in the second layer is obtained by inverting the Laplace transform using the Post approximation (Post, 1930). After presenting a delta function as input to the first layer, the activity of units in the second layer resembles the firing rates of scale-invariant time cells. The model can be implemented as a two layer feedfoward neural network where the weights can be explicitly computed as a function of the time constants of the nodes in the first layer. Here we give a brief overview of the mathematical model and emphasize the connection to our neural circuit that will be introduced later.

The goal of this method is to reconstruct a vector-valued function over the time leading up to the present f(*t′* < *t*). For simplicity we focus our attention on a single component *f* (*t*). As shown in Figure 3a, this input stimulus is fed in parallel into a series of leaky integrators *F*(*s,t*):

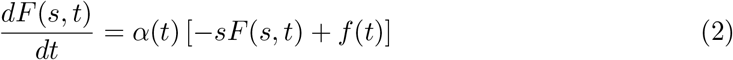

Here *F*(*s,t*) is the activity of the node labeled by *s*. α(*t*) is an an externally controlled parameter. For now, we assume that α(*t*) = 1.^1^ We can observe from Equation 2 that the set of activities *F*(*s, t*) are just the Laplace transform of the original stimuli with Laplace index *s*.

**Figure 3.**
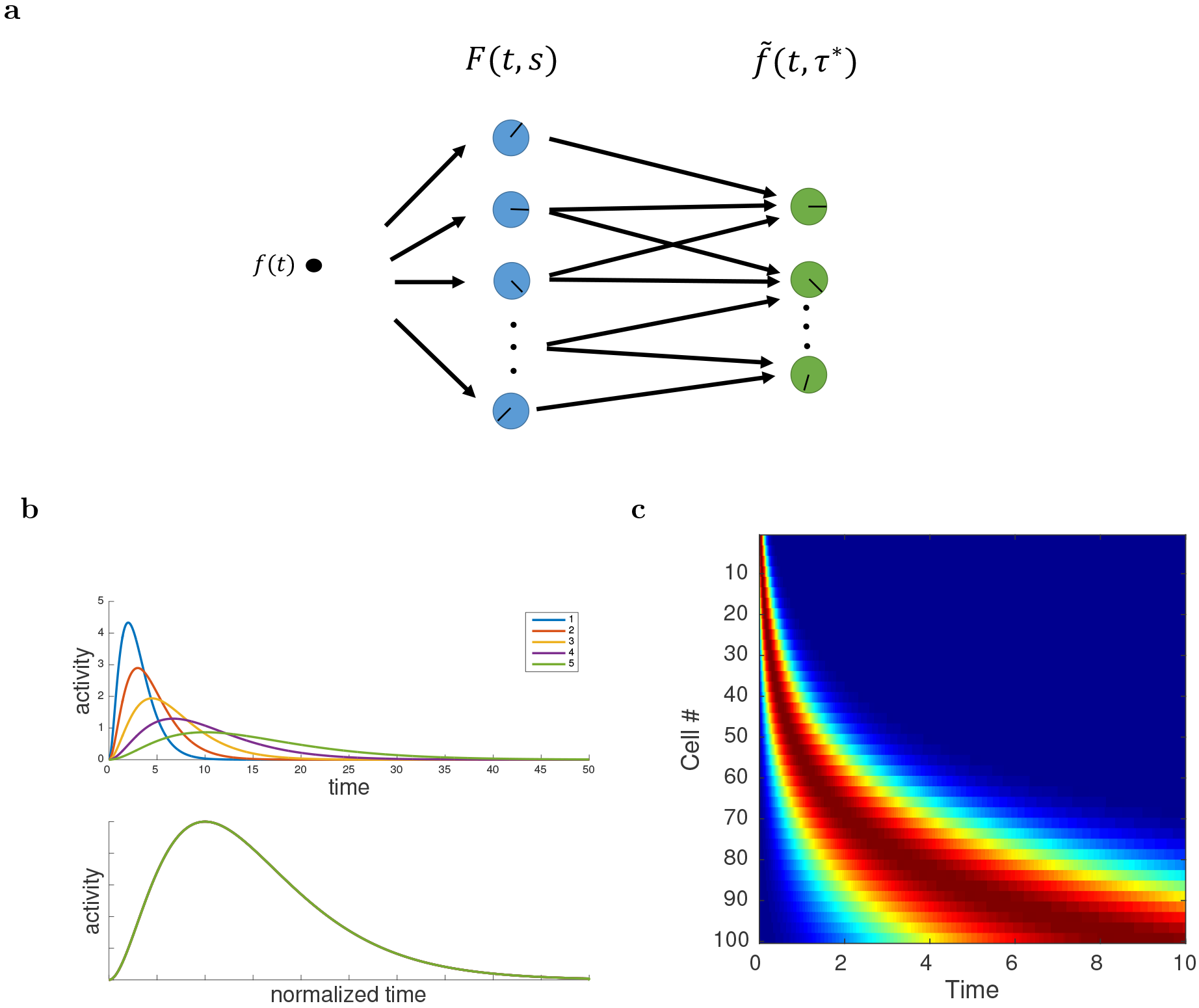
Illustration of the scale-invariant mathematical model for sequentially-activated time cells. **a.** Rather than a simple chain, in this formulation, an input is provided in parallel to a set of leaky integrators, *F*. These units provide a feedforward input to another set of units 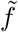 that function like time cells. The direction of the ‘clock hands’ within the neurons indicate the time constant of that neuron (peak time for sequentially-activated cells and decay time constant for leaky integrators) **b.** The activation of 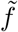 nodes after a delta function input in the mathematical model is scale-invariant. **Top:** The activation of 5 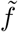 nodes with different time constants 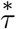 (Equation 7). *k* = 2 is chosen in the inverse Laplace transform. **Bottom:** The same functions rescaled by 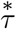 along the x axis and by the maximum activation along y axis as in Figure 2. Unlike the chaining model, the five lines exactly overlap each other, showing the scale-invariant property, **c.** Heatmap from the mathematical model. The time at which a time cell activates is ultimately controlled by the time constant of the integrators that provide input to it. Choosing the time constants of the leaky integrators controls the number density of time cells.

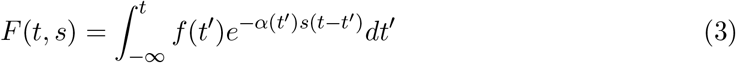

More specifically if we take the stimulus to be a delta function *f*(*t*) = δ(0), the neuron represented by the *F* node will simply have an exponentially decay firing rate with a decay time constant of 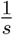

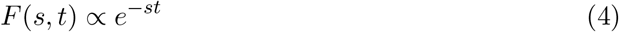

The activity of the *F* nodes is transformed by a second layer of 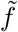 nodes. The 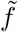 nodes are in one-to-one correspondence with the *F* nodes. At each moment the activity of the *F* node labeled by *s* is transformed in the following way:

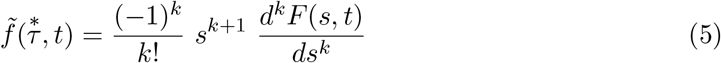

where 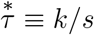 is a parameter that indexes the nodes in 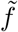^2^ and *k* is some integer that we later identify to be related to the precision of the inverse Laplace transformation. The only time-varying part in Eq. 5 is *F*. It will turn out that the value of 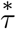 specifies the time that each unit in 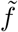 has its peak activation following a delta function input.

The above transformation is an inverse Laplace transform in the sense that
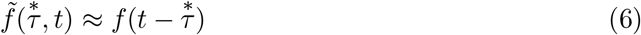
where 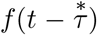 is the value of the stimulus function a time 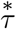 prior to the present. The approximation becomes exact when *k* → ∞ (Post, 1930). Because there are many units in 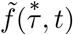 that set of units traces out the past values of the input function such that an approximation of the entire function is available at time *t*. Thus we can see that the set of activations 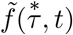 constitutes a faithful representation of the original stimulus function delayed by 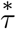. This is true regardless of the form of the function *f*(*t*).

To better understand the properties of 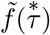, consider the form it has if *f*(*t*) is a delta function at time zero. Then, each node in *F*(*s*) decays exponentially as *e^-st^* and each 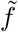 node is given by:
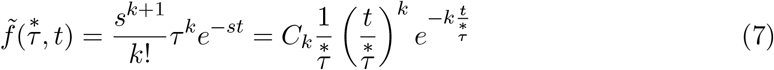
where *C*_*k*_ is a constant that depends only on the choice of *k* and we have substituted 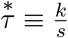 in the last step.

Equation 7 has properties that resemble the firing rate of a time cell with a peak firing rate at 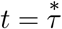. Note that the time-dependence of this expression depends only on the fraction 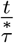. Thus if we rescale the x axis according to 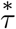, and the y axis by the maximum activity, the firing activity of all the cells will coincide, as shown in Figure 3b. Thus, the activity of nodes in 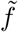 is scale-invariant.

This mathematical framework produces a set of functions that resembles the firing rates of time cells. Moreover this mechanism gives rise to time cells that are scale-invariant, which would be a desirable property for the brain to possess. However, it is not clear whether it is possible for neural circuits in the brain to actually implement this hypothesized mechanism. We will demonstrate that this is indeed neurally realistic by constructing a biologically detailed neural circuit that utilizes a biophysical model of exponentially decaying persistent firing neuron (Tiganj et al., 2015) to perform the computation of this mathematical model, thereby generating a set of scale-invariant time cells.

The values of 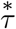 in 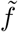 are controlled by the values of *s* in *F*. It remains to specify the distribution of values of *s* and thus 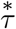. In order to preserve scale-invariance, equate the information content of adjacent nodes (Shankar & Howard, 2013) and enable 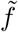 to implement Weber-Fechner scaling (Howard & Shankar, 2018), we choose the values of 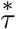 to be logarithmically spaces as shown in Figure 3c. This is equivalent to choosing the number density of *s* to go down like *s*^-1^. Power law distributions of time scales emerge in physical systems under quite general circumstances (e.g., Amir, Oreg, & Imry, 2012)

### A biophysical model implementing this mathematical framework

The mathematical framework requires two physical processes. One is a set of exponentially-decaying cells with a spectrum of time constants. For this we follow (Tiganj et al., 2015) with a set of integrate-and-fire neurons equipped with a slowly decaying calcium-dependent non-specific cation current (Fransén et al., 2006). The second process is an implementation of the operator to approximately invert the Laplace transform in Eq. 5. We will implement this with a neural circuit with realistic synaptic conductances. First, however, we discuss how to implement the derivatives Eq. 5 with discrete values of *s*.

### The weight matrix for inverse Laplace transform W_*L*_

By Equation 5, the connection weights should depend on the discretized *k*th derivative with respect to *s*. To write derivatives in matrix form, imagine there are three successive *F* nodes, with labels *s*_–1_, *s*_0_ and *s*_1_. Note that the first derivative of a function with respect to *s*_0_ can be approximated as a weighted average of the slope of the line connecting the successive points on the curve.

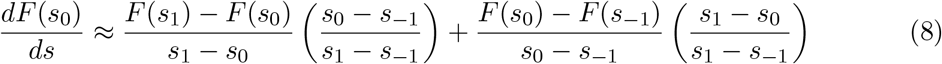

The factors in the parentheses account for the fact that the accuracy of the slope further away from the point *s*_0_ is a less accurate estimate of the derivative.
 [^3^ This approximation works when the second derivative 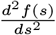 does not change sign along the interval[*s*_–1_,*s*_1_]. This is always the case for *F*(*s, t*).]

The derivative as expressed in Equation 8 is a linear combination of *F*(*s*_1_), *F*(*s*_0_) and *F*(*s*_–1_) with coefficients determined by *s*_1_, *s*_0_ and *s*_–1_. Therefore if we represent the function *F*(*s*) by a discretized vector of values

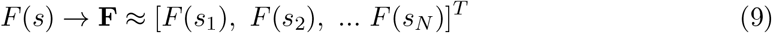

we can write the first derivative as a matrix,

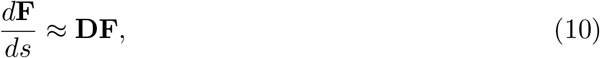

where the matrix D is given by

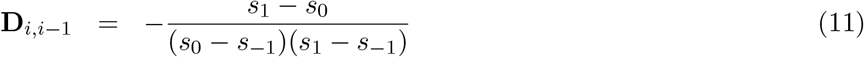

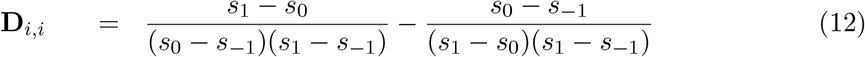

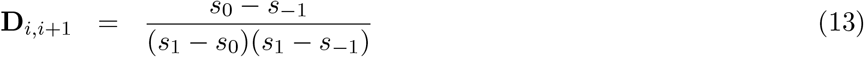

for *i* = 1,2,…,*N* and all the other elements are 0. The *k*th derivative can be obtained by simply multiplying *k* of these matrices.

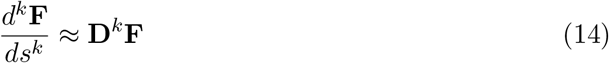

After we have the matrix representation of the kth derivative, the activity for a given 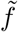 node 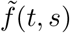 can be approximated by a linear combination of the *F* node activities *F*(*s,t*).

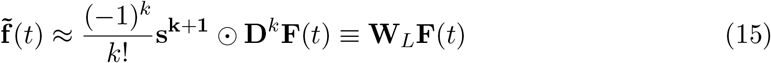

where ⊙ represents element-wise multiplication, and 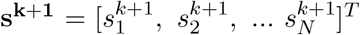. Thus the inverse Laplace transform is readily implemented in a neural network via a weight matrix **W**_*L*_.

In our simulation we will choose *k* = 2, so for a given 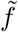 node labeled by *s*_0_ its activity at any given time *t* will depend on its five nearest neighboring *F* nodes, *F(t,s_–2_), F(t,s_-1_), F(t,*s*_0_), F(t,s_1_)* and *F(t,s_2_).* In this simulation, we have 9 *F* nodes labeled by *s*_1_ to *s*_9_. According to above they will generate 5 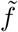 nodes.

**Figure 4.**
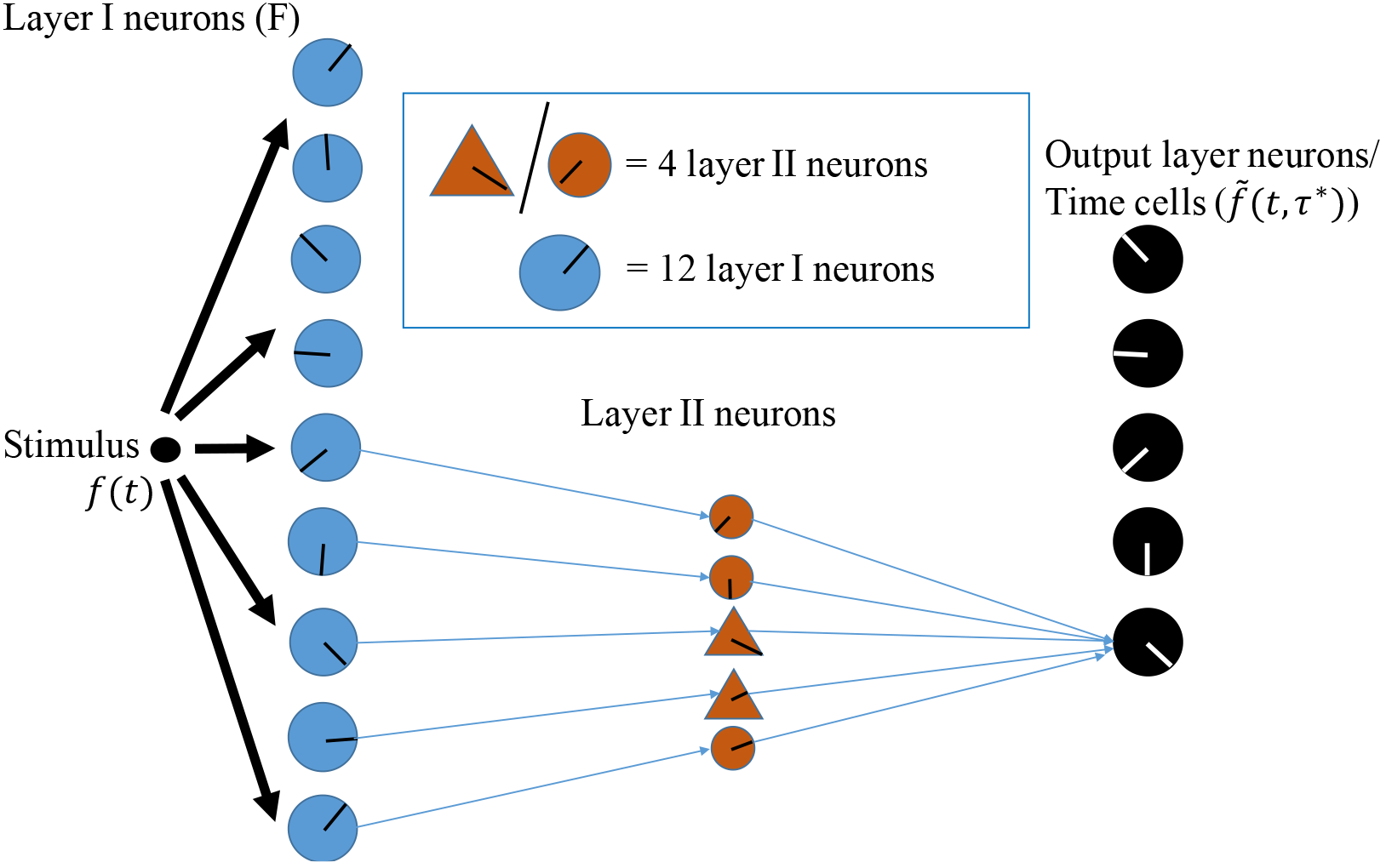
Schematic of the architecture of the biological network. Only a subset of the layer II neurons are drawn. Each blue circle represents 12 layer I neurons. Each orange circle represents 4 excitatory layer II neurons. Each orange triangle represents 4 inhibitory layer II neurons. Each black circle represents one output layer neuron. The direction of the ‘clock hands’ within the neurons indicates the time constant of that neuron. In the actual simulation, 9 groups of 12 persistent spiking layer I neurons connect to 5 output layer neurons (later identified as time cells) via weights generated by a matrix representation of the inverse Laplace transform **W**_*L*_. In order to satisfy Dale’s Law, an excitatory/inhibitory layer II neuron is placed on each positive/negative connection. In total, there are 108 layer I neurons, 100 layer II neurons and 9 output layer neurons.

### A biophysical model for exponentially decaying persistent firing neurons

Biologically, time cell sequences have been observed stretching out to at least a minute. Because 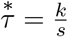 implementing these equations requires some mechanism implementing time constants of at least tens of seconds. It is non-trivial to identify a biophysical mechanism that can result in persistent spiking that lasts over that period of time. While recurrent connections with appropriate eigenvalues would implement this property perfectly well, here we follow previous work that uses known single-cell properties to build long time constants. Tiganj et al. (2015) developed a computational model of single neurons that uses a calcium-activated nonspecific (CAN) cationic current to achieve decay time constants up to several tens of seconds under realistic choice of parameters. Here we utilize that same model as the neural realization of the *F* nodes.

The model works because cells contain a slowly-deactivating current that depends on calcium concentration. Because this current causes spikes, and because spikes cause an influx of calcium, this mechanism can result in very long functional time constants. Because the functional time constant depends on spiking, mechanisms that alter the amount of current needed to cause a spike also alter the functional time constant.

The dynamics of the model are summarized as follows:

1. During the interspike interval, the membrane potential *v_m_(t)* is modeled to be only affected by the CAN current
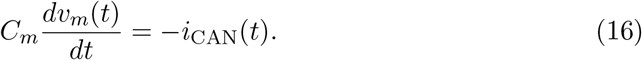

where *C*_*m*_ is the membrane capacitance and *i*_*cAN*_ is the CAN current.
II. The CAN current is given by
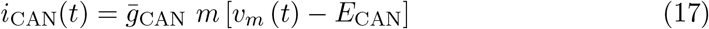

Here 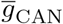 is the maximal value of the ion conductance measured in 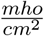, *E*_*CAN*_ is thereversal potential of the CAN current ion channels and *m* is a dimensionless quantity between 0 and 1 that is associated with the activation of the CAN current ion channels.
III. Critically, the value of *m* depends on the concentration of calcium. Following previous computational work (Traub, Wong, Miles, & Michelson, 1991; Fransén, Alonso, &Hasselmo, 2002), we let the activation parameter for the ion channel *m* change from moment to moment according to

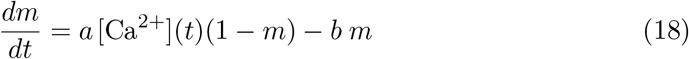

where *a* and *b* are free parameters. Following Fransén et al. (2002), we choose *a* and*b* as 0.02 and 1, respectively, in this simulation.
IV. The dynamics of the calcium concentration is given by an exponential decay during the interspike interval.

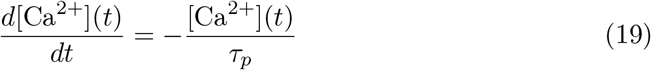

where *τ*_*p*_ is the decay time constant.
V. Critically, a fixed amount influx of calcium whenever the cell fires an action potential.

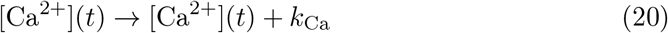

Tiganj et al. (2015) showed both analytically and numerically that under the appropriate assumptions the firing rate will be approximately exponentially decaying with a functional decay time constant τ that far exceeds the decay time constant τ_p_ of the calcium concentration from Eq. 19:

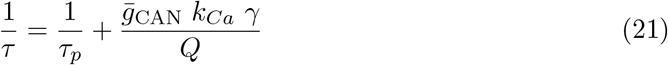

where 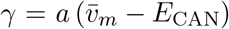, 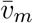 is the average membrane potential during a spike and Q is the total charge influx during a spike. Note that this expression allows infinite values of the functional time constant. This expression holds when the neurons are in the linear regime and when the change of *m* is much faster than the change of calcium i.e., 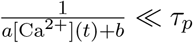 and the interspike interval is much less than *τ_p_,* the decay time constant of the calcium concentration. In the simulation we choose the parameters to satisfy the two conditions above, so that the firing rates are well approximated by exponential decaying functions with a broad range of time constants.

### Layer I neurons implementing *F* nodes

Layer I consists of 108 integrate-and-fire neurons driven by the CAN current described above, modeling the *F* nodes from the mathematical model. The set of model neurons spans 9 different values of *s* with 12 neurons for each value. To model the experimental finding of logarithmically-compressed timeline, the time constants 1/*s* were chosen to be logarithmically spaced between 2 s and 83 s by adjusting maximum CAN current conductance 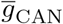 and initial calcium concentration in the CAN driven persistent firing neuron model above according to Table 1. All the model parameters are summarized in Table 1

**Table 1.** List of the parameters of the model

| Variable | Label | Value |
| --- | --- | --- |
| layer I Neurons |  |  |
| threshold potential | $v_t$ | -40mV |
| reset potential | $v_{reset}$ | -70mV |
| reversal potential of CAN current | $e_{CAN}$ | -20mV |
| capacitance | $C_m$ | $10^{-4} \mu F$ |
| calcium decay constant | $\tau_p$ | 1000ms |
| dynamical parameter for m | a | 0.02 |
| another dynamical parameter for m | b | 1 |
| maximum CAN current conductance | $g_{CAN}$ | 0.0023, 0.0031, 0.0036, 0.0039, 0.0042, 0.0043, 0.0044, 0.0045, 0.0045mho/cm<sup>2</sup> |
| calcium influx every spike | $k_{Ca}$ | 0.001 (calcium concentration is defined to be unitless) |
| initial calcium concentration | $Ca(0)$ | 0.05/0.0376/0.0322/0.0294/0.0278/0.0268/0.0262/0.0258/0.0255 |
| layer II Neurons |  |  |
| time constant | $\tau_i$ | 250ms |
| threshold potential | $v_{t,i}$ | -50mV |
| reset potential | $v_{reset,i}$ | -50.2 mV |
| output layer Neurons |  |  |
| time constant | $\tau_{post}$ | 50ms |
| threshold potential | $v_{t,post}$ | -50mV |
| reset potential | $v_{reset,post}$ | -50.2 mV |
| Synaptic Potential (modeled as alpha function) |  |  |
| time constant for NMDA receptor | $\tau_{alpha,NMDA}$ | 45ms |
| time constant for GABA<sub>A</sub> receptor | $\tau_{alpha,GABA}$ | 15ms |
| duration of a single synaptic potential |  | 300ms |

### Layer II neurons relay the activity and ensure Dale’s law

The expression for the synaptic weights **W**_L_ in Eq. 15 requires both positive and negative connections. The 9 groups of layer I neurons are connected to 5 output layer neurons (time cells) via connection matrix **W**_*L*_ with *k* = 2. If there were only direct connections between layer I neurons and the output layer neurons, implementing **W**_*L*_ would violate Dale’s Law. To ensure adherence to Dale’s law we place a layer II neuron on every connection. The layer II neurons are inhibitory or excitatory depending on the sign of the connection weight that they convey to the output neurons, as shown in Figure 4.

To keep the firing rates of the layer II neurons in a reasonable regime, the PSPs (postsynaptic potentials) coming from 3 layer I neurons from the same group are used as the input to one layer II neuron. Layer II neurons are modeled as leaky integrate and fire neurons. The time evolution of the layer II neurons is given by

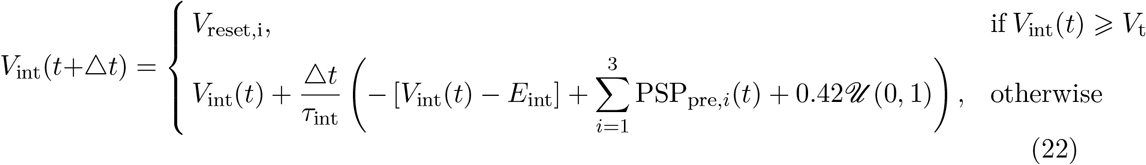

where *E*_int_ is the resting potential for the layer II neurons, τ_int_ is the membrane time constant, PSP_pre,*i*_ is the PSP coming from layer I neuron *i* and a noise term 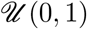 represents a random number drawn uniformly from (0,1). We added a background current of 0.42mA so that the layer II neurons are in their linear regime, an essential point to ensure scale-invariance.

### Layer II cells project to output layer cells

PSPs generated by the layer II neurons with 5 different time constants of the intermediate neurons are summed up and provide input to the output layer neurons, as in Figure 4b. The weight matrix **W**_*L*_ is implemented by different PSP amplitudes of individual layer II neurons, which can be computed from Equations 11–15. We also rescaled the individual PSPs so that the inputs to the different output layer neurons have the same maximum. This ensures that the output layer neurons are all in their linear regimes. A biophysical mechanisms to achieve such regime could be due to homeostatic synaptic scaling in which the activity of neurons regulates the magnitude of synaptic inputs (Turrigiano, Leslie, Desai, Rutherford, & Nelson, 1998). Later we will elaborate that as the time constants of nearby neurons become closer (i.e., *s*_*i*+1_ – *s*_*i*_ → 0), the receptive field will closely resemble an off-surround, on-center one.

### Output layer cells model the 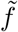 nodes in the mathematical framework

The output layer neurons are also leaky integrate and fire neurons.

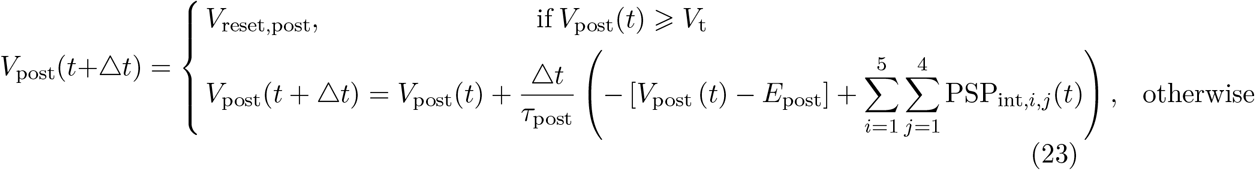

where *i* indexes different time constants and *j* indexes the different layer II neurons with the same time constant.

Although CAN current is prevalent in pyramidal neurons, the amount of persistent spiking in different populations of entorhinal cortex neurons varies, with less persistent spiking in stellate cells (Klink & Alonso, 1997). Thus, the CAN current may be present in different magnitudes in different neuronal populations. We have used simple integrate and fire neurons in the output layer as a simplified initial representation, but future implementations could include a more complex range of membrane currents.

We use alpha functions for modeling the synaptic potentials and Euler’s method with a time step of 0.1 ms in MATLAB 2016a to implement differential equations.

## Results

There are two primary results in this paper. The first is that the simulated output layer neurons, like time cells, fire sequentially in response to a delta function input and the sequential firing is approximately scale-invariant. This property comes from receptive fields that can be understood as off-center/on-surround in the projections to the output layer units. The second is that the simulated neural sequence can be rescaled by adjusting the gain of the layer I neurons. Before describing those results, we first describe the activity profile of each of the layers in turn.

### Layer I neurons showed exponential decay with a range of time constants

The layer I neurons as shown in the bottom layer of Figure 4 are driven by the input to the network. They are equipped with an internal CAN current and have persistent, exponentially decaying firing rates. The decay time constants are logarithmically spaced between 2.04 s and 83.49 s. This is achieved by adjusting the maximum CAN current conductance *GCAN* and initial calcium concentration according to Table 1.

The dotted lines are exponential functions; the degree to which the firing rates align with these theoretical functions confirm that the firing rates indeed decay exponentially. This is in accordance with the activity of the *F* nodes in the mathematical model.

There are 9 groups of layer I neurons in total, each neuron within a group has the same time constant. Within each group there are 12 layer I neurons with the same parameters. For every 3 of them, their PSPs are summed up and sent as input to one layer II neuron, as shown in Equation 22 and Figure 4.

### Layer II cells also decayed exponentially

The layer II cells shown in the middle layer of Figure 4 are leaky integrate and fire neurons, with parameters given in Table 1. Driven by upstream neurons with exponentially decaying firing rates, they also display firing rates that decay exponentially, at least initially. At very long times they maintain a background firing rate of around 1 Hz due to the background current as described in Equation 22. This ensures that the layer II neurons are always in the linear regime, which is crucial for exact scale-invariance of the computation performed by this neural circuit.

**Figure 5.**
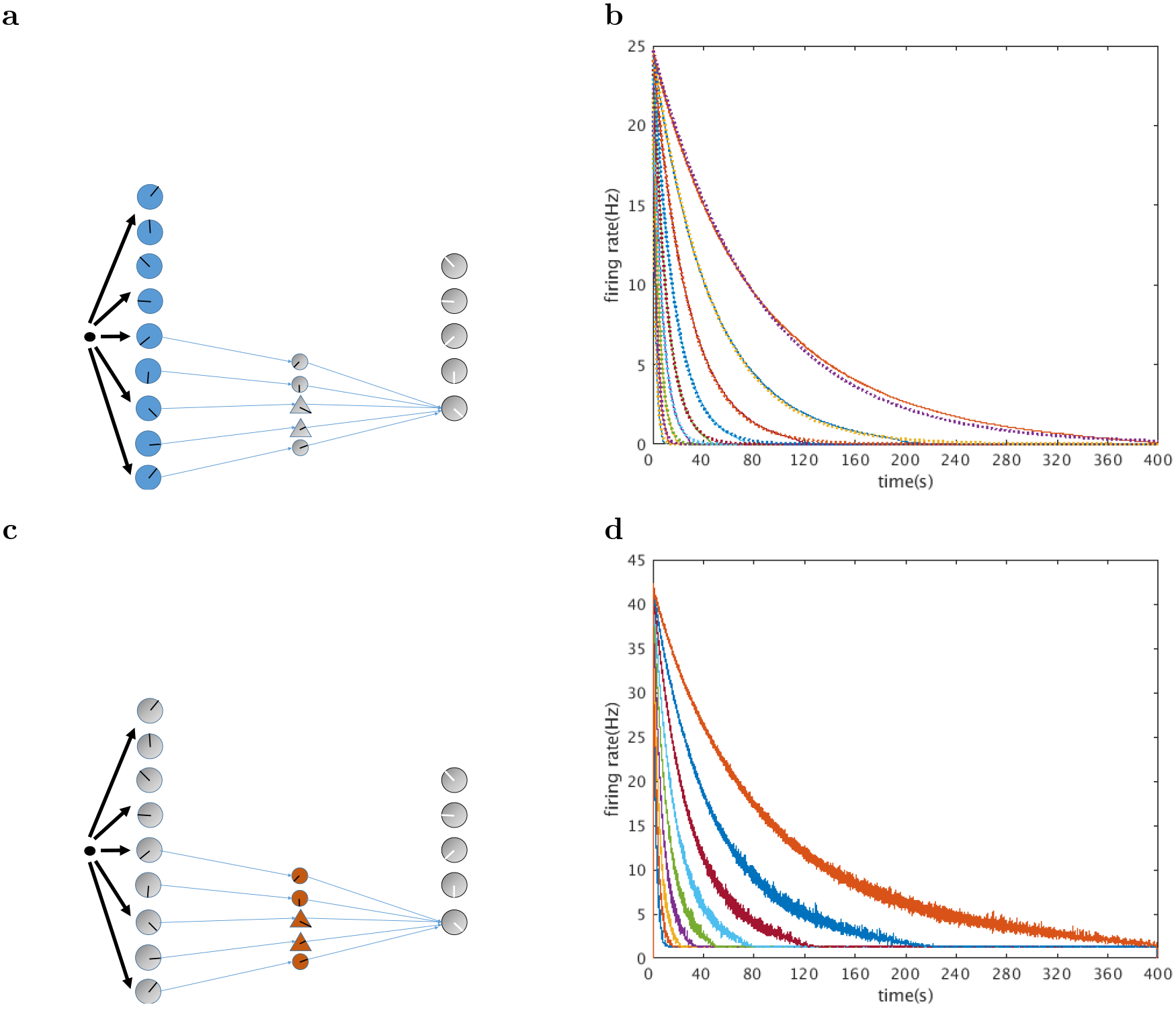
Neurons in layer I and layer II decay exponentially. **a.** Layer I is composed of exponentially decaying persistent spiking neurons with CAN current. These cells are driven by the network input, **b.** Firing rates for layer I neurons. Nine example neurons with time constants logarithmically spaced between 2.04 s and 83.49 s are shown, **c.** Layer II is composed of leaky integrate and fire neurons. They are driven directly by the downstream persistent spiking layer I neurons. They serve the role of exciting and inhibiting the neural activity from layer I to the output layer in order to complete the computation of the inverse Laplace transform, **d.** Firing rates for 9 layer II neurons with different decay constants. All firing rates are averaged over 100 trials.

The PSPs of the layer II neurons contribute differently to the output layer neurons due to the different amplitude of their individual PSPs. The PSPs of layer II neurons with 5 different time constants provide input to one output layer neuron, as shown in Figure 4. Since each layer II neuron is only involved in one connection, Dale’s Law is satisfied by simply choosing the layer II neuron to be excitatory if it corresponds to a positive weight, and vice versa. The model does not have constraints on the specific type of synaptic transmitter used. For this particular simulation, the excitatory neurons have NMDA receptors with a time constant of 45 ms (Otis & Mody, 1992) and the inhibitory neurons have *GABA*_*A*_ receptors with a time constant of 15 ms (Perouansky & Yaari, 1993).

### Post-synaptic neurons fired sequentially and were approximately scale-invariant

The post-synaptic cells shown in the top layer of Figure 4 are also modeled as leaky integrate and fire neurons. When driven by the PSPs from the layer II neurons, their firing rates resemble the mathematical expression of Equation 7. As shown in Figure 6 their peak firing times scale with the width of their firing fields. When rescaled according to peak firing times, their firing rates overlap with each other. This indicates that the output layer neurons fire sequentially and have a firing rate profile that is time-scale invariant.

### The weight matrix **W**_*L*_ approximates an off-center, on-surround receptive field when time constants are densely spaced

In the above simulation we chose a specific series of time constants and the neighboring index *k* = 2 in the inverse Laplace transform, and the connectivity pattern is given by Equation 15. For any choice of time constants and *k* value, the general form of connectivity can be derived from the matrix **W**_*L*_ in Equation 15. We found that as the time constants of nearby 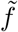 nodes become closer(i.e., *s*_*i*+1_ – *s*_*i*_ → 0), the receptive fields will become more similar to a symmetrical off-center, on-surround one. To this end, we ran an additional simulation with 99 (instead of 9) time constants logarithmically spaced between 2 s and 50 s. The results are shown in Figure 7. The receptive fields for all the time cells have the same off-surround, on-center shape, and their firing rates are still scale-invariant. In vision, receptive fields like this can be learned from natural scene statistics by maximizing statistical independence (Bell & Sejnowski, 1997) or sparsity (Olshausen & Field, 1996). If temporal and visual information processing reflect similar principles (Howard, 2018), it is possible that the receptive field in our model could also reflect the adaptation of our mechanism of temporal information processing to statistical properties of the world.

**Figure 6.**
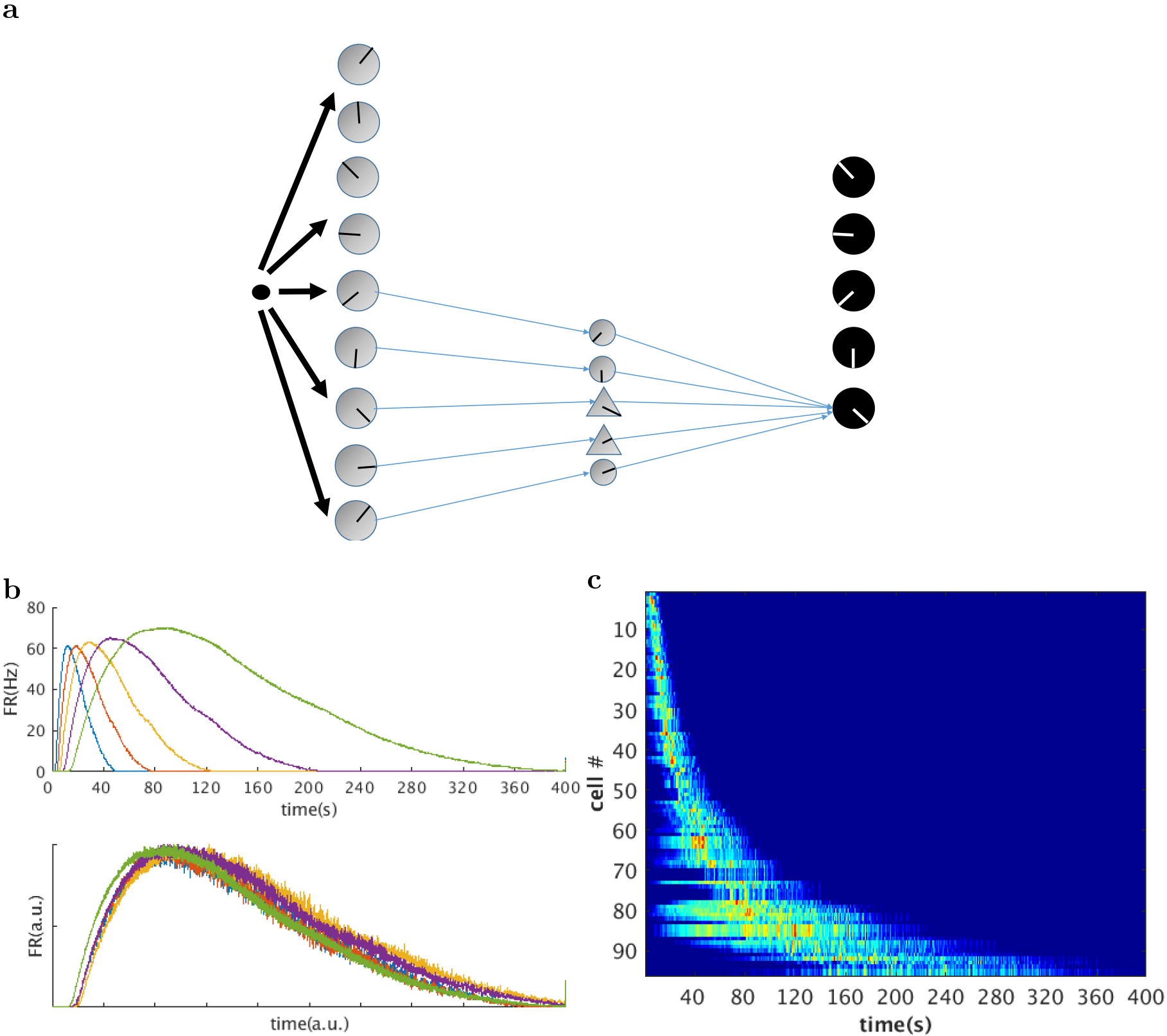
Our model produces scale-invariant time cells that resemble neural data. **a.** The output layer consists of 5 leaky integrate and fire neurons which we identify as time cells, **b.** From top to bottom: output layer firing rates for the five time cells generated by this network. Rescaled version of the firing rates for the five neurons. It is clear that the firing rates coincide with each other when rescaled, showing that the firing rates for the time cells are indeed scale-invariant. Firing rates are averaged over 100 trials, **c.** Heatmap generated from 100 simulated time cells. Compare to Figures 1, 2c, and 3c.

### The network exhibits linearity in response to a square wave input

In the above simulation we presumed that the time cells fire during a delay period, whose start is signaled by some stimulus which we abstracted as a delta function. In reality we would be interested in the response of the network to a temporally extended stimulus. To this end we ran the model with a square wave input that lasted for 150s. Since the Laplace transform of a continuous function is just the convolution of that function with the Laplace transform of a delta function, the response of the time cells in theory would just be the convolution of the stimulus with the impulse response given by Equation 7. As shown in Figure 8, the firing rates of the layer I and layer II neurons faithfully represent the Laplace transform of the square wave input, and the firing rates of the time cells agree with the theoretical prediction (shown as solid lines in the figures).

### Time rescaling of time cells can be achieved by globally changing f-I curves of layer I neurons

This mathematical approach can readily simulate the “time rescaling” phenomenon, where the firing fields of time cells are rescaled by the length of the delay interval (MacDonald et al., 2011; Mello et al., 2015; Wang, Narain, Hosseini, & Jazayeri, 2018). In recurrent neural network models, time rescaling manifests itself as a collective phenomenon where the neural trajectory sweeps through similar space but with different linear (Hardy, Goudar, Romero-Sosa, & Buonomano, 2017; Wang et al., 2018) or angular(Goudar & Buonomano, 2017) speeds. In previous work based on reservoir computing frameworks, rescaling requires learning new sets of weights. On the contrary in the current framework, rescaling of the neural sequence is achieved simply by cortical gain control, i.e., a global change in the slope of the f-I curves among all the layer I neurons.

Figure 9 shows the results of two simulations where the speed of the sequence was rescaled by 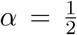 and α = 2, according to Equation 2. This is equivalent to the time constants of all the layer I neurons being rescaled by 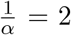 and 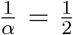 respectively. Notice that according to Equation 21, the time constant is controlled by the maximum CAN current conductance 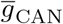, calcium influx after a spike *k*_*Ca*_ and the charge required for a spike Q. These variables all affect the slope of the f-I curves of the layer I neurons. Here we altered the conductance 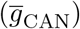 of all the layer I neurons. This potentially reflects the effect of acetylcholine on the activation of the muscarinic receptors. Previous study has shown that activation of the muscarinic receptors activates a calcium-sensitive, nonspecific cation current (Shalinsky, Magistretti, Ma, & Alonso, 2002) which induces persistent firing (Hasselmo & McGaughy, 2004). High levels of acetylcholine are also associated with attention, which is related to the change in cortical gain (summarized in (Thiele & Bellgrove, 2018)). We focused on changes in CAN current conductance because of this data on the effect of acetylcholine. Less data is available on modulations of other physiological parameters.

**Figure 7.**
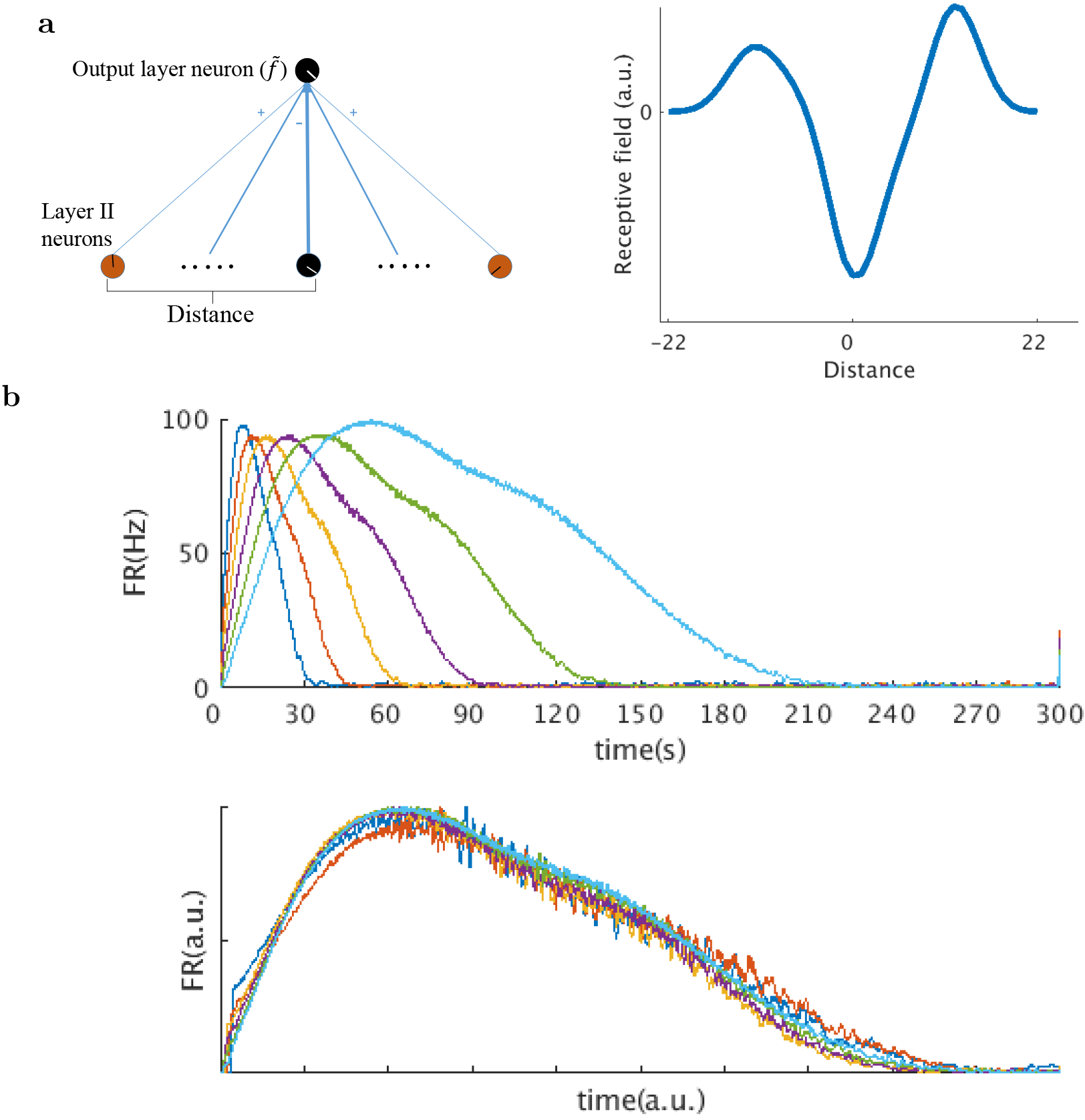
Increasing the number of layer I neurons result in off-center, on-surround receptive fields for time cells. We showed that as the nearby time constants get closer, the shape of the receptive field becomes similar to an off-center, on-surround one. We simulated the network activity using an augmented version of the model with 99 time constants for layer I neurons spanning 2s-50s. The resulting receptive fields become almost off-center, on-surround (**a** right, distance indicates the distance from the layer II neuron with the same label *s* as the output layer neuron (black circles, **a** left), line thickness in **a** left schematically represents the amplitude of the receptive fields). The firing rates of the time cells still remain scale-invariant, **(b** top, activity of 6 representative time cells, **b** bottom: the firing rates rescaled along the x axis by peak time.)

**Figure 8.**
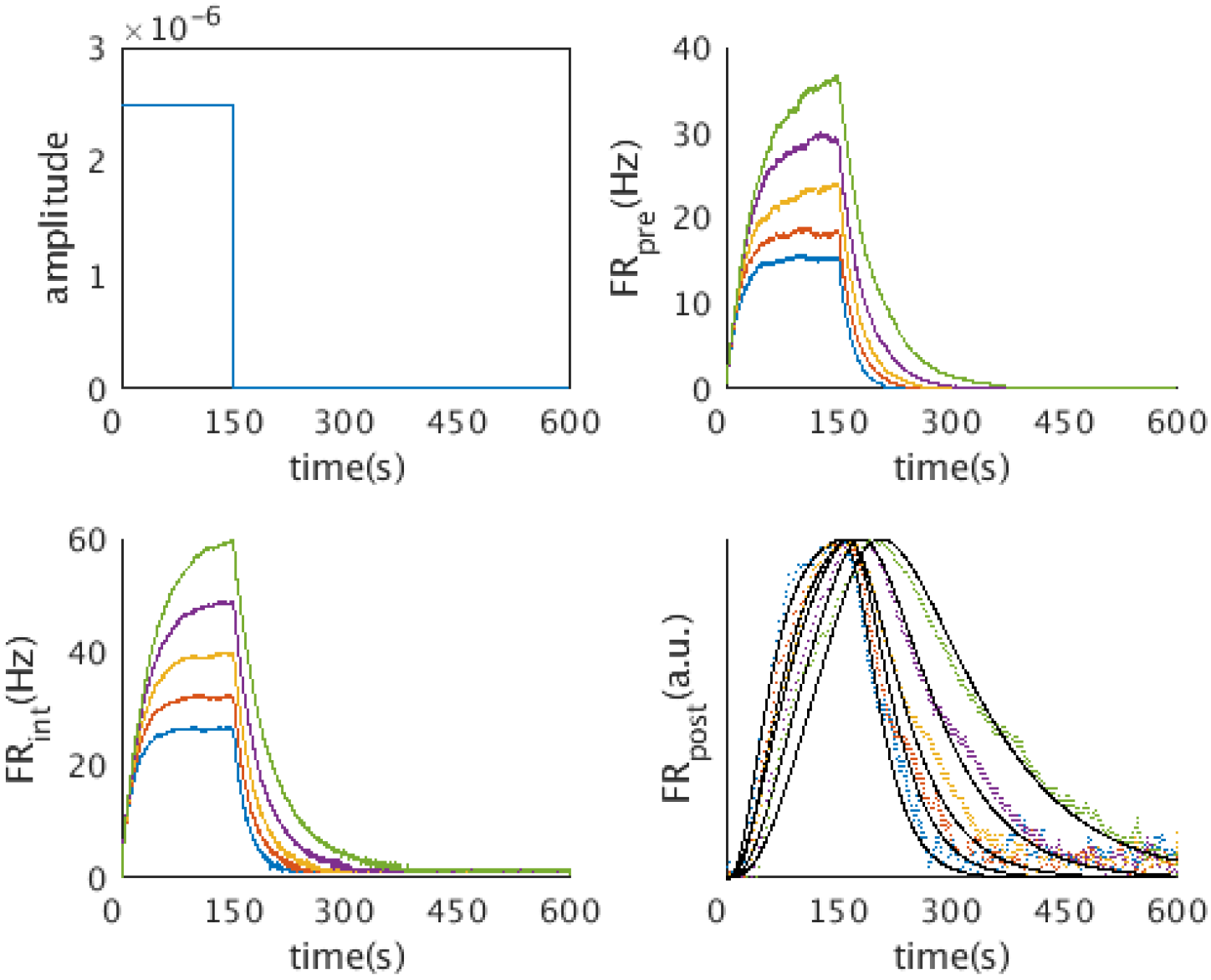
The network implements the Laplace transform and inverse for a temporally extended stimulus. We simulated the network activity with a square wave input (top left). The network activity agrees with the theoretical prediction from the mathematical framework. The layer I neurons and the layer II neurons exhibit the activity of a charging capacitor (5 representative layer I neurons, top right; 5 representative layer II neurons, bottom left). The firing rates of time cells (bottom right, 5 representative time cells shown) agree with the prediction from the mathematical framework (black line)

Indeed as shown in Figure 9a,b changing the slope of the f-I curves changes the time constants of layer I neurons. Figure 9c shows the firing rates of the same time cells before and after the change in α Their firing fields appear rescaled by the scaling factor 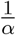
, in accordance with observation(MacDonald et al., 2011; Mello et al., 2015).

Figure 9b shows the peak times of 55 time cells before and after remapping. The sequentially activated time cells follows a straight line, indicating that the time cells indeed code relative time during an interval. [^4^ Note that in this simulation we used an augmented version ofthe model where the layer I neurons span 99 time constants.]

## Discussion

We proposed a neural circuit that encodes the Laplace transform of an input function and then approximately inverts the Laplace transform to produce a series of sequentially firing cells. The time constants and the peak firing times of the time cells range from a few seconds to almost 100 seconds. Critically, the firing rates of the sequentially-activated time cells are scale-invariant. This provides a possible neural substrate for the scalar timing behavior observed across a wide range of timescales in behavioral tasks, and also approximates the neurophysiological recordings of sequentially activated cells that have been observed across a wide range of regions of the cortex including the hippocampus(MacDonald et al., 2011; Salz et al., 2016), PFC (Tiganj et al., 2017) and striatum (Adler et al., 2012; Mello et al., 2015).

## Constraints on the neural circuit to preserve scale-invariance

A biological constraint on the neural circuit that can cause deviation from scale-invariance is the input-output function (f-I curve) of the layer II neuron. Only when the layer II neurons are in their linear regimes can they faithfully relay the temporal information from the presynatic neurons to the output layer neurons. Since we modeled the layer II neurons as leaky integrate and fire neurons, their f-I curves are discontinuous near the threshold input value. Thus some background firing is required for the layer II neurons to be in their linear regimes. Also some steady background firing for the layer I neurons would not change the scale-invariance property, since a constant shift in *F(s,t)* would not affect the derivative that contributes to the inverse Laplace transform.

Alternatively, any type-I model neuron with a linear f-I curve would satisfy the biological constraint imposed by scale-invariance, with or without background firing. By appropriately modeling an adaptation current, a log-type f-I curve could be transformed into a linear one (Ermentrout, 1998).

**Figure 9.**
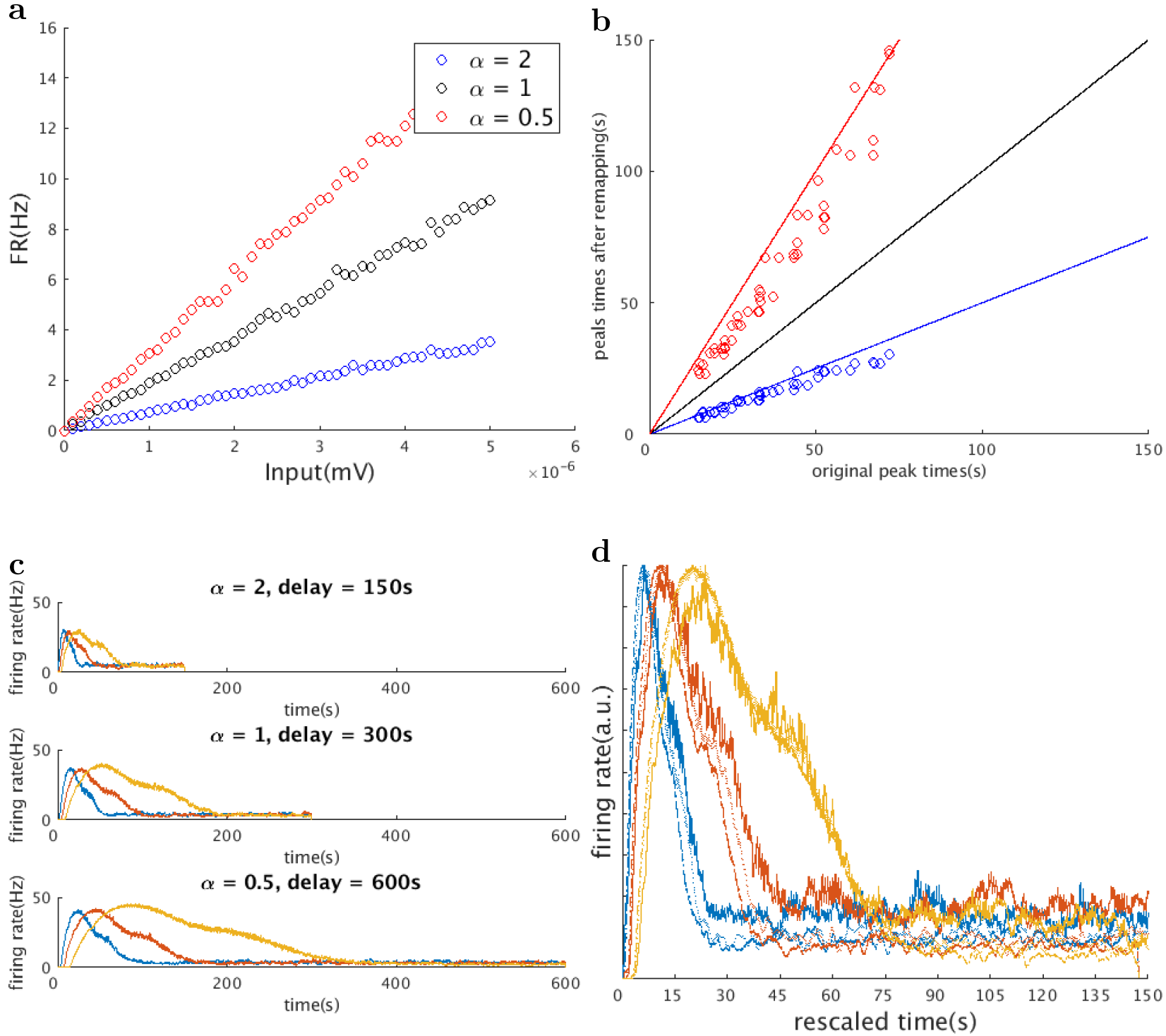
Rescaling of time cell firing rates by a global change of the slope ofthe f-I curves. We simulated the “time rescaling” phenomenon described in MacDonald et al., 2011, where the firing fields of time cells are rescaled according to the length of the delay interval. We globally changed the slopes of the f-I curves of all the layer I cells by means of changing the maximum CAN current conductance 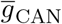(**a**, three f-I curves of the same layer I cells when the delay interval is changed to half (blue), twice (red), and the same as the original). This results in the rescaling of the peak times of all the time cells (**b**, red: peak times become twice the original, blue: peak times become half the original. Three straight lines indicate *y* = 0.5*x*(blue), *y* = *x*(black) and *y* = 2*x*(red)). Firing rates of 3 representative time cells are shown in c (before remapping (**c**,top), when the delay interval is changed to one half(**c**,middle) and twice (**c**,bottom) of the original, same color indicates the same cell). The firing rates of the same time cell under different delay lengths coincide when the time axis is rescaled according to the peak times (**d**). Color indicates the same cell in (**c**), thick line: α = 2 dotted line: α = 1, thin line: α = 0.5)

Additional biological features of this model are the use of decaying persistent spiking activity, which resembles the persistent firing properties observed in intracellular recordings from slice preparations of the entorhinal cortex (Klink & Alonso, 1997; Tahvildari, Fransén, Alonso, & Hasselmo, 2007; Jochems, Reboreda, Hasselmo, & Yoshida, 2013; Egorov et al., 2002) and peririnal cortex (Navaroli et al., 2011). Mechanisms of decaying persistent firing has also been observed in other structures such as hippocampus (Knauer, Jochems, Valero-Aracama, & Yoshida, 2013) and prefrontal cortex (HajDahmane & Andrade, 1996). In addtion, there are in vivo recordings showing a spectrum of timescales across cortex (Bernacchia, Seo, Lee, & Wang, 2011; Murray et al., 2014; Meister & Buffalo, 2017; Tsao et al., 2017).

## Alternative approaches to implementing the mathematical model

There are other neural circuit models that produce sequentially activated neurons, but to our knowledge, the present model is the first one that has the additional feature of scale-invariant neuronal activity. However, functionally identical models with different biological realizations of the same equations might also be possible. For example, rather than implementing long functional time constants *via* intrinsic currents, one could construct an analog of Eq. 2 using recurrent connections. For example, Gavornik and Shouval showed that in a spiking recurrent neural network trained to encode specific time intervals, units exhibit persistent spiking activity (Gavornik & Shouval, 2011). Other neural circuits for computing the inverse are also possible. The computation of the inverse Laplace transform amounts to a suitable linear combination of inputs from cells with exponentially decaying firing rates. Poirazi, Brannon, and Mel (2003) showed, in a detailed compartmental model of a hippocampal CA1 pyramidal cell, that the dendritic tree functions as a two-layer artificial neural network. Thus a single dendritic tree could implement something like **W**_*L*_ used here. It is also within the realm of possibility that the exponentially decaying firing rates could be replaced with slow dendritic conductances.

## Acknowledgments

We thank two reviewers for helpful comments. This work is supported by ONR MURI N00014-16-1-2832 and NIBIB R01EB022864.

## Footnotes

1 By modulating α(*t*) with velocity, the model can produce place cells (Howard et al., 2014).

2 Note that this definition of 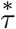 differs from the notation in some previous papers where 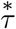 was defined to be negative. We adopt this convention for convenience here.

